# Diet-responsive proteogenomic effects following short-term restriction of animal products in humans

**DOI:** 10.1101/2025.07.21.665884

**Authors:** Alexandros Simistiras, Ozvan Bocher, Christina Emmanouil, Anargyros Skoulakis, Stavros Glentis, Nikolaos Scarmeas, Eleftheria Zeggini, Konstantinos Rouskas, Antigone S Dimas

## Abstract

The effect of diet on genetic regulation in humans remains largely unexplored. Here, we investigated gene-by-diet interactions in a unique group of apparently healthy individuals (N=200) who alternate between omnivory and dietary restriction of animal products for religious reasons. Using longitudinal plasma proteomic and genotypic data, we identified diet-responsive cis-protein quantitative trait loci (cis-pQTLs) including gain of genetic regulatory effects for LBR and MSRA, proteins linked to cholesterol and methionine metabolism respectively. LBR-associated diet-responsive cis-pQTL rs74148404 showed suggestive evidence of colocalization with obesity exclusively under dietary restriction, suggesting diet-dependent differential genetic risk for disease. Additionally, a novel dietary restriction-associated cis-pQTL for metabolic regulator FGF21 colocalized with eosinophil and platelet traits pointing to diet-sensitive immunometabolic signalling. By parallel profiling of a continuously omnivorous control group (N=211), we also uncovered seasonally dynamic gain and loss of genetic regulation for proteins linked to apoptosis in immune system pathways (MAVS, CASP3, PDLIM7 IL12RB1), effects likely masked by animal product restriction. These findings reveal dynamic diet- and season-sensitive regulatory mechanisms with implications for precision nutrition and individualized disease prevention strategies, and underscore the need to integrate environmental context into genetic studies of health and disease.

## Main

Most studies exploring the impact of regulatory variants in humans capture effects of steady-state environmental conditions. Steady-state expression quantitative trait loci (eQTLs) however, explain at most ∼40-50% of GWAS associations, and a fraction of unexplained associations likely stems from gene-environment interactions^1,2^. Understanding the complex interplay between environmental factors, such as dietary intake, and genetics has the potential to enrich our understanding of key mechanisms that contribute to higher level phenotypes, including disease. Diet is a key determinant of the metabolic environment in which our genes function and constitutes a modifiable risk factor for diseases including obesity, cardiovascular disease, type 2 diabetes, and cancer^3,4^. To date, dietary restriction studies, involving interventions that restrict intake of total energy or of specific macronutrients, have highlighted molecular pathways sensitive to nutritional modulation^3,5,6^. However, little is known about how our genes respond to changes in dietary intake. Given the largely untapped potential of dietary modifications to ameliorate health, there is a growing need to elucidate interactions between genetic variants and diet in order to better understand diet-responsive molecular mechanisms driving effects on health.

In genetically diverse mice, response to dietary restriction was shown to be a complex, polygenic trait driving highly individualized effects on health and lifespan^7^. In baboons, a recent study identified genetic regulatory effects on RNA levels that emerged in response to a prolonged high-cholesterol, high-fat diet^8^. Diet-responsive cis-eQTLs were found to be enriched in human GWAS genes associated with metabolic phenotypes, including BMI, cholesterol, atherosclerosis and liver cirrhosis. In humans, a study identified extensive effects of complete caloric restriction on plasma protein levels and showed that the abundance of these proteins can be genetically regulated^5^. However, direct effects of diet on genetic regulation of molecular profiles remain largely underexplored to date.

In the present study, we explored the impact of diet on proteogenomic regulation by profiling a unique group of apparently healthy individuals who alternate between omnivory and dietary restriction of animal products for religious reasons, in a consistent and highly structured pattern involving restriction for 180-200 days annually. Collectively, our results demonstrate diet-responsive gain and loss of genetic regulatory effects for proteins with key roles for cellular homeostasis. Furthermore, we have connected diet-responsive regulatory variants to higher level health-related phenotypes and highlight proteins through which dietary restriction likely modulates disease risk. The insights derived from this study enhance our understanding of the molecular mechanisms through which diet affects health, of the population-level variability that exists for health-related responses to dietary intake, but also aid the development of therapeutic approaches that mimic beneficial effects on health exerted by specific dietary patterns.

## Results

### Population sample

We profiled a unique group of apparently healthy individuals from Greece (FastBio study: religious <u>Fast</u>ing <u>Bio</u>logy) who alternate between omnivory and dietary restriction of animal products for religious reasons (periodically restricted PR group, N=200) (**Fig. 1**, **Supplementary Table 1**). This study population and the dietary pattern adhered to has been described in detail elsewhere^9,10^. Briefly, PR individuals abstain from meat, fish, dairy products and eggs for 180-200 days annually, in a consistent and highly structured pattern involving four extended periods of restriction throughout the year, and restriction on Wednesdays and Fridays of each week. The consistency of this dietary pattern and the predictability of the shift between omnivory and animal product restriction render this study comparable to a dietary intervention study. We compared findings to a continuously omnivorous control group of individuals profiled in parallel (non-restricted, NR group, N=211). Participants were profiled at two time points: T1 in autumn, covering a period of omnivory for both dietary groups, and T2 in early spring, covering a three-to-four-week period of restriction for PR individuals, during Lent. Our study design therefore comprises four different contexts as dietary group by time point combinations (PR at T1, PR at T2, NR at T1, NR at T2) **(Fig. 1)**. Typically, during Lent, PR individuals undergo restriction of protein (as a proportion of total energy intake), driven chiefly by abstinence from almost all sources of animal protein, which is not accompanied by a decrease in total energy intake^11,12^. We have previously shown that during this period PR individuals undergo extensive metabolic reprogramming, with mostly beneficial effects on health^9,10^, which encompasses reductions in plasma levels of cholesterol and other lipid classes, of branched-chain amino acids, and altered abundance of key immunometabolic proteins, including potent metabolic hormone FGF21^9^. When considering genetic ancestry, FastBio participants clustered close to individuals of Tuscan, Cypriot, Bulgarian, and Turkish descent (**Extended Data Fig. 1**).

**Figure 1.**
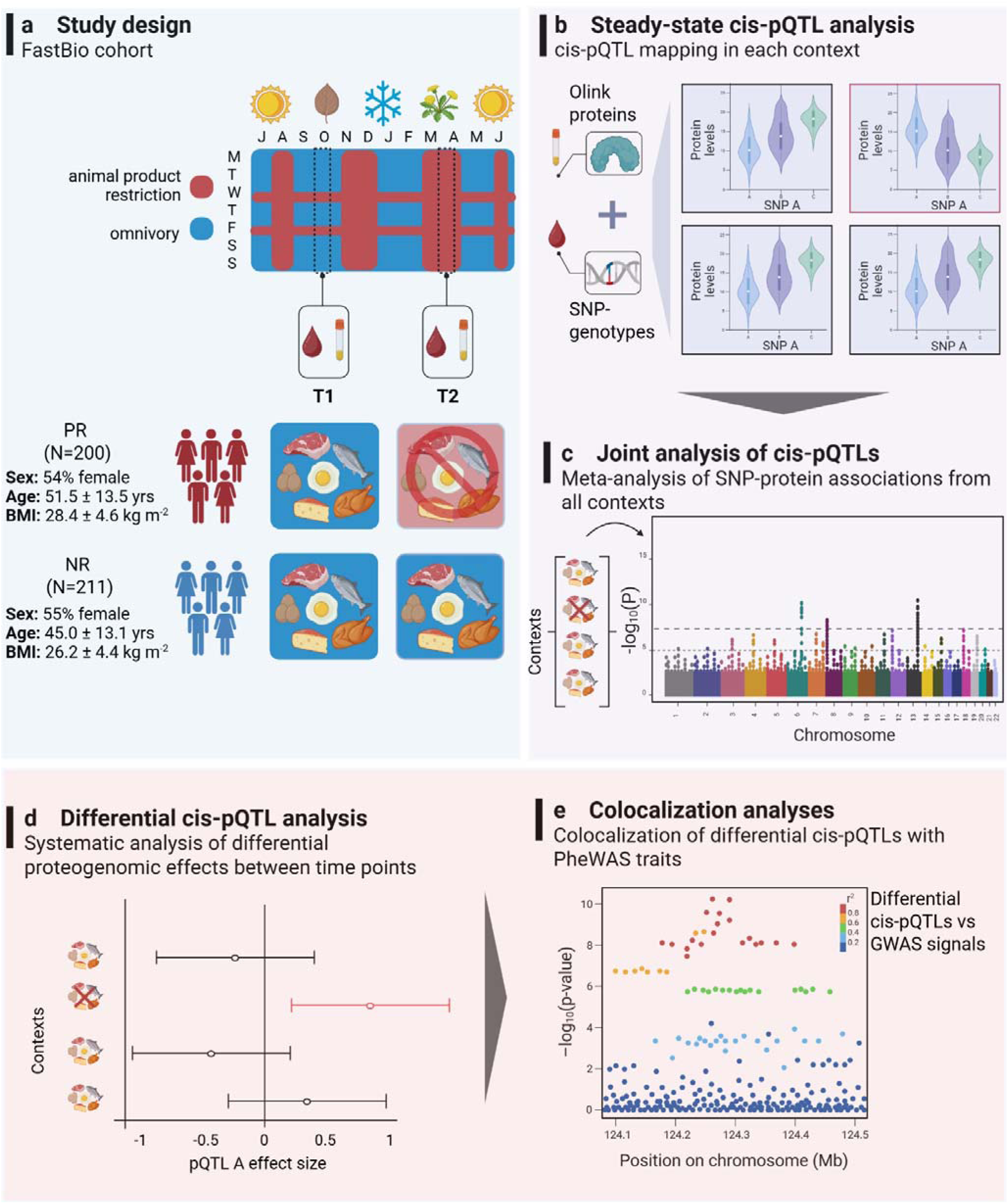
Study overview. **a,** FastBio study design and participant demographics. **b,** Steady-state cis-pQTL mapping for each time point by dietary group combination (context). **c,** Joint analysis of steady-state cis-pQTLs from all contexts using mashr. **d,** Systematic analysis to define differential cis-pQTLs between sampling time points for each dietary group based on mashr results. **e,** Colocalization analysis for differential cis-pQTLs with PheWAS-selected traits. Created using bioRender. PR: periodically restricted group; NR: non-restricted group; T1: time point 1; T2: time point 2.

### Map of steady-state cis-pQTLs

We mapped steady-state cis protein QTLs (cis-pQTLs) for each diet by time point combination (context), and identified 585, 511, 559 and 550 independent cis-pQTLs for PR at T1, PR at T2, NR at T1 and at T2 respectively, and an average of 431 proteins associated with at least one cis-pQTL (pGenes) per context (**Extended Data Table 1, Supplementary Table 2**). Collectively across contexts we identified 630 pGenes with 32% possessing more than one independent cis-pQTL (min = 2; max = 5). The vast majority of cis-pQTLs were shared between contexts, with concordant direction of effect, and clustered around the TSS of their pGene (**Extended Data Fig. 2a-c**). Eighty four percent of top cis-pQTLs from all contexts (k = 630) were replicated in the UK Biobank pharma proteomics project (UKB-PPP)^13^ at a Bonferroni-corrected threshold of 7.94e-5 (**Extended Data Fig. 2c**). Forty four percent of top cis-pQTLs mapped in intronic regions, while 16% and 14% were found in upstream and downstream gene regions respectively (**Extended Data Fig. 2d**). Over half (59.4%) of the top cis-pQTLs mapped on known regulatory regions, including transcription factor binding sites, or overlapped with eQTLs (**Supplementary Table 3**). Top cis-pQTLs explained a median of 10.1% (IQR = 13%) of the variance in protein abundance (**Supplementary Table 4**). For 22 proteins the top cis-pQTL explained over 50% of the variance, underscoring the presence of strong cis-regulatory effects. For example, rs72620873, a cis-pQTL shared between all contexts, explained ∼ 59% of the variance in levels of SIRPA, an immune receptor that maintains self-tolerance and is linked to cancer, transplant rejection, autoimmunity, and neurodegeneration when dysregulated^14^.

### Differential cis-pQTL mapping and colocalization with GWAS traits

To identify cis-pQTLs with diet-responsive effects, we first used mashr in a meta-analysis framework to jointly analyse SNP-protein associations from all contexts detected through steady-state mapping. In total, we identified 583 cis-pQTLs (lfsr < 0.05), 80.3% of which were also found through steady-state mapping. Next, we tested for cis-pQTLs exhibiting differential effects between time points for both dietary groups and although close to 99% of cis-pQTLs were shared between all contexts, we detected nine variants displaying strong differential effects, involving gain or loss of genetic regulatory effects, from T1 to T2 (**Fig. 2**).

**Figure 2.**
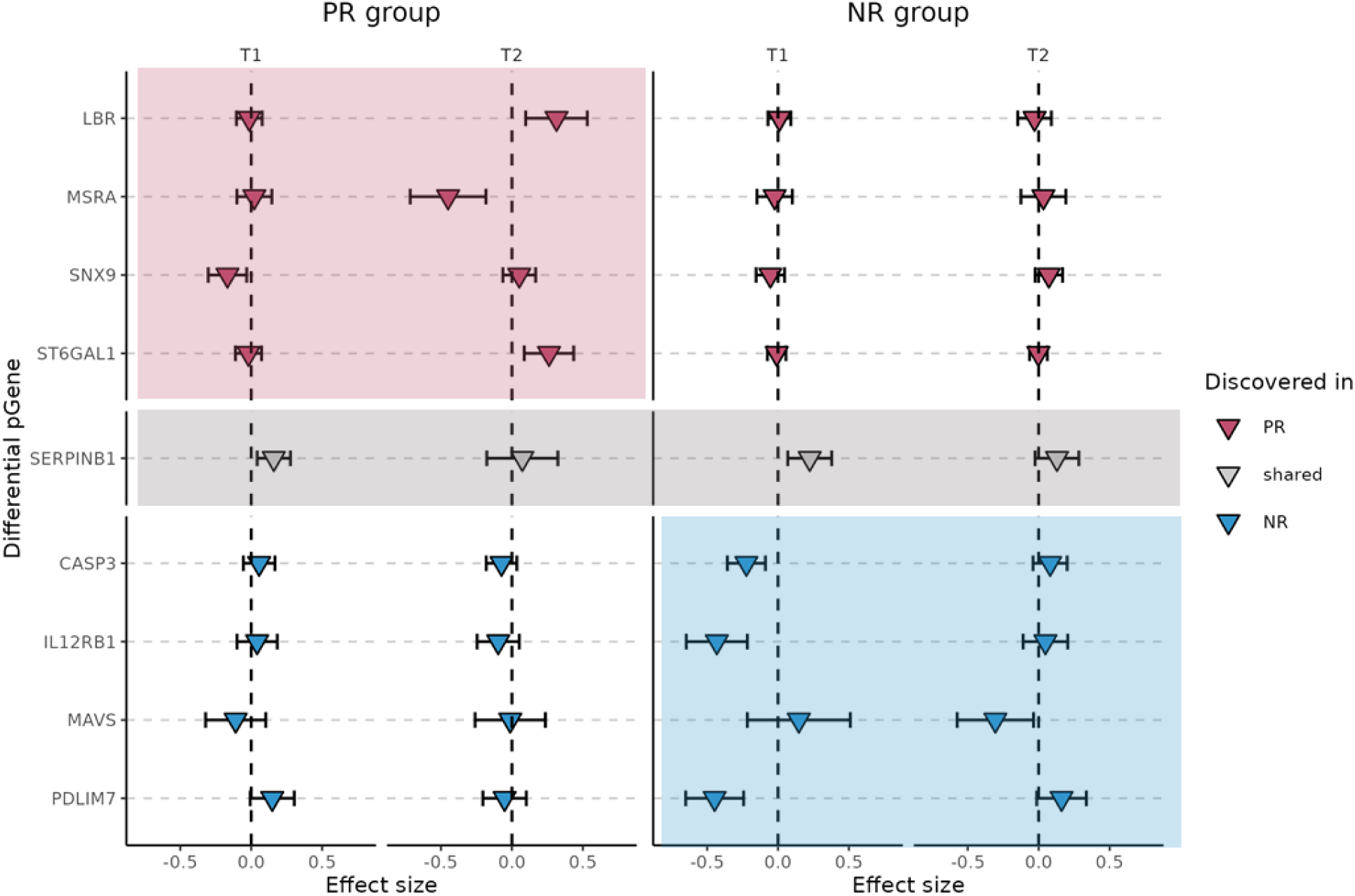
Differential cis-pQTLs detected from T1 to T2 and corresponding pGenes. Forest plot depicting the effect size of the association between each SNP-protein pair with differential effects at each time point and for each dietary group. Triangles show SNP effect sizes (posterior mean estimate from mashr) and the error bar represents +/- 2 posterior standard deviations of the effect size. Red: differential cis-pQTLs detected in PR; grey: differential cis-pQTL detected in both dietary groups; blue: differential cis-pQTLs detected in NR (control group). Shaded boxes indicate the dietary group in which differential cis-pQTL was detected. PR: periodically restricted group; NR: non-restricted group; T1: time point 1; T2: time point 2.

We found four previously unreported instances of diet-responsive gain (LBR, MSRA, ST6GAL1) or loss (SNX9) of genetic regulatory effects on protein abundance (**Fig. 2**). We also uncovered loss of genetic regulatory effects at T2 for SERPINB1 in both dietary groups, whereas in the control group, we detected gain of genetic regulatory effect at T2 for a single protein (MAVS) and three instances of loss of regulatory effects (PDLIM7, CASP3, IL12RB1). Seven out of nine differential cis-pQTLs mapped at distances between 270-993 kb from the TSS, in a pattern similar to that detected for diet-responsive cis-eQTLs in baboons^8^ (**Extended Data Fig. 2b**). Furthermore, this subset of cis-pQTLs was not found in UKB-PPP^13^, suggesting the possible existence of previously unreported, environmentally-driven genetic effects that may not be captured by cross-sectional studies. On the contrary, the differential cis-pQTLs that mapped close to their genes TSS (rs2293772-SERPINB1; rs11668601-IL12RB1) were replicated in UKB-PPP and may capture broader effects induced by environmental fluctuations that are experienced by the general population, such as seasonality.

Next, we explored whether differential cis-pQTLs contribute to higher level health-related phenotypes using colocalization analysis, and compared findings between contexts. Through phenome wide association (PheWAS) queries we found that six out of nine differential cis-pQTLs, or their proxies (r^2^ > 0.8), had at least one reported GWAS association at p-value < 1e-5 (**Supplementary Table 5**). Through single trait colocalization of GWAS traits with differential cis-pQTLs, we found suggestive evidence of colocalization (PPH4 = 0.55) of diet-responsive locus LBR with obesity. Furthermore, for loci detected only in the control group, we found evidence of colocalization (PPH4 > 0.8) for IL12RB1 with gamma glutamyl transpeptidase (GGT) and 11 other traits, for CASP3 with intrinsic epigenetic age acceleration, and for PDLIM7 with arm fat percentage (**Extended Data Table 2**). Given that we had a limited set of potential instrumental variables (IVs), we were not able to conduct Mendelian randomization (MR) analyses to estimate causal effects between proteins and health outcomes.

### Gain of genetic regulatory effect on LBR levels in response to reduction in dietary intake of cholesterol

We uncovered a previously unknown regulatory variant that emerged only during dietary restriction of animal products and was associated with the abundance of inner nuclear membrane protein LBR (rs74148404; lfsr = 0.014; in PR at T2) (**Fig. 3a**). LBR interacts with chromatin and lamins, has an essential role in endogenous cholesterol biosynthesis, and is required for viability of human cell lines under cholesterol starvation^15^. This genotype-dependent upregulation of LBR may be a compensatory response to reduced dietary intake of cholesterol during animal product restriction compared to omnivory^9,16^. We validated diet-responsive genetic effects of rs74148404 on plasma levels of multiple cholesterol-containing lipoproteins in PR individuals (e.g. total cholesterol p.adjusted = 0.057 PR at T1; p.adjusted = 0.024 PR at T2) and found no effects in the control group (**Fig. 3b, Supplementary Table 6**).

**Figure 3.**
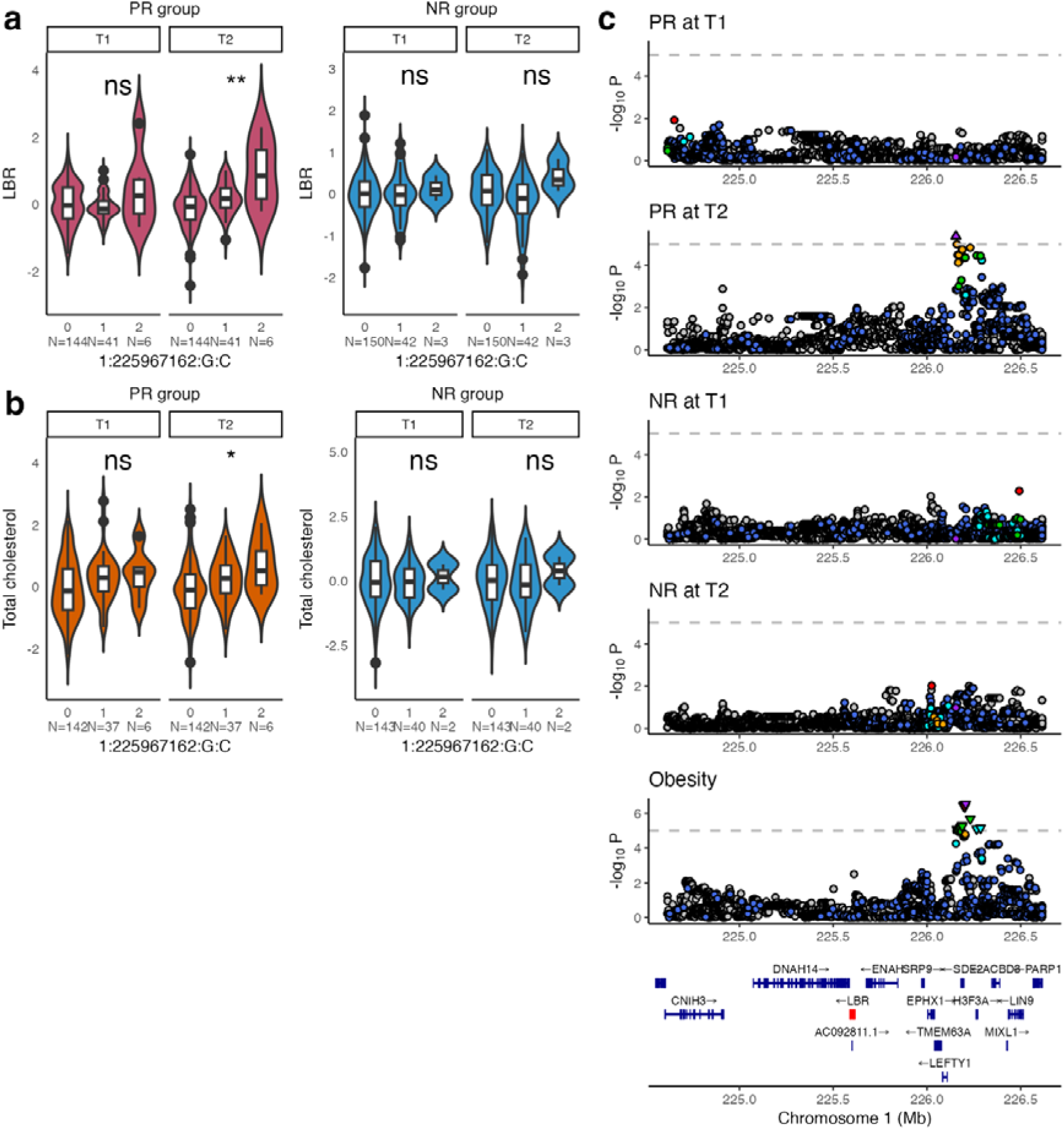
Diet-responsive cis-pQTL rs74148404 is associated with LBR levels and colocalizes with obesity only during animal product restriction. **a,** Dosage effect of differential cis-pQTL 1:225967162:G:C (rs74148404) on LBR levels in PR and NR groups. **b,** Validation of rs74148404 genetic effects on total cholesterol levels in PR individuals during animal product restriction (T2), no effect was detected in the NR group. **c,** Stacked regional plots for colocalization results of rs74148404 and obesity in each context; rs74148404 has suggestive evidence of colocalization with obesity only during dietary restriction (PR at T2, PPH4 = 0.55) and in no other context (PPH4 range: 0.01 to 0.06 in other contexts). Asterisks in panel **a** indicate significance as *: p.adjusted < 0.05; **: p.adjusted < 0.01; ***: p.adjusted < 0.001 (p.adjusted based on 1,000 permutations from steady-state cis-pQTL mapping); ns = non-significant. Asterisks in panel **b** indicate significance as *: p.adjusted < 0.05; **: p.adjusted < 0.01; ***: p.adjusted < 0.001 (p.adjusted based on FDR = 5%); ns = non-significant. PR: periodically restricted group; NR: non-restricted group; T1: time point 1; T2: time point 2; p.adjusted = adjusted p-value.

Notably, we found suggestive evidence of colocalization for rs74148404 with obesity, only during animal product restriction (PPH4 = 0.551 in PR at T2) (**Fig. 3c**). Although we could not assess the causal effect of LBR on obesity through MR, we note that the LBR-increasing C allele at rs74148404 was associated with a marginal protective effect against obesity (beta = −0.001; p = 5.5e-05) raising the possibility that carriers may be slightly more protected against obesity through the action of LBR, when practicing dietary restriction. This finding may contribute to our understanding of why individuals who follow similar dietary patterns may exhibit differential risk for obesity^17^ and provides broader insight into mechanisms shaping diet-dependent differences in disease risk.

Moreover, our previous findings revealed that LBR abundance decreased at T2 in the control group (log_2_FC = −0.097; p = 1.65e-6)^9^. When considering non-carriers of the diet-responsive C allele at rs74148404 from the PR group (N = 144), we also found decreased abundance of LBR at T2 (logLJFC = –0.070; p = 0.003) (**Supplementary** Fig. 1). This suggests that under animal product restriction, carriers of the diet-responsive allele drive LBR levels in the opposite direction of changes occurring under omnivory. Given known seasonal fluctuations in cholesterol levels^18^, we speculate that abundance of LBR may also exhibit seasonal variation that could be counteracted by dietary restriction of animal products.

Adding to the idea of diet-responsive genetic effects, rs74148404, which is located 539 kb downstream of the LBR TSS, maps on the distal end of a predicted RXRA::VDR motif. These motifs regulate the transcription of genes in response to vitamin D, a nutrient whose intake is also typically lower in plant-based diets^16^. Vitamin D and cholesterol metabolism is interconnected^19^, but to date little is known about the underlying regulatory mechanisms. Finally, given the role of LBR in tethering peripheral heterochromatin to the nuclear envelope^20^, our findings may extend beyond cholesterol homeostasis pointing to potential diet-dependent genetic effects on chromatin organization. Further exploration of these discoveries may enhance our understanding of mechanisms underlying cholesterol homeostasis.

### Genotype-dependent downregulation of MSRA in response to reduction in dietary intake of methionine

We found a previously unreported instance of diet-responsive gain of genetic regulatory effects on MSRA levels (rs74891397; lfsr = 0.006 in PR at T2) (**Fig. 4a**). MSRA is an enzyme that repairs oxidative damage by reducing both protein-bound and free oxidized methionine residues, thereby restoring protein function and biological activity^21^. Plant-based diets are associated with lower levels of oxidative stress relative to omnivorous diets^4^ and typically involve lower intake of dietary methionine^22,23^. We suggest that the diet-dependent genetic effect on MSRA levels occurs in response to reduced methionine intake and may reflect a genotype-specific lower requirement for MSRA-mediated repair during dietary restriction. To explore genetic effects of rs74891397 further, we measured plasma levels of methionine in a subset of PR individuals (n = 60; 27 C allele carriers at rs74891397 and 33 non-carriers) and found that diet-responsive genetic effects extended to methionine levels (linear regression model; beta = 1.949; se = 0.782; p = 0.016 in PR at T2) (**Fig. 4b**).

**Figure 4.**
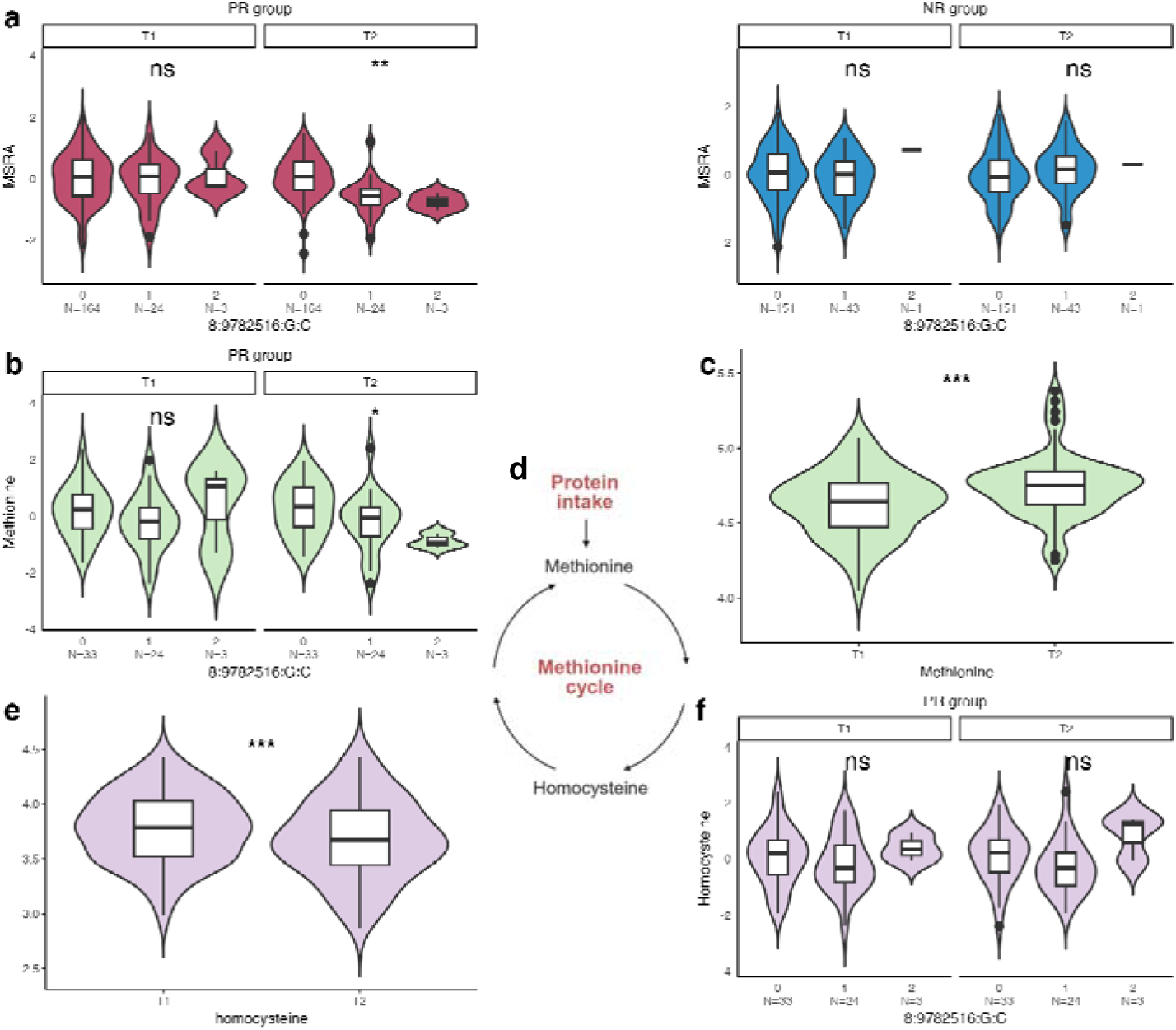
Diet-responsive cis-pQTL rs74891397 associated with MSRA levels. **a,** Dosage effect of differential cis-pQTL 8:9782516:G:C (rs74891397) on MSRA levels in PR and NR groups. **b,** Validation of rs74891397 genetic effects on plasma methionine levels in PR individuals (n = 60) during animal product restriction. **c,** Distribution of methionine levels in PR individuals (n = 60) at both time points. **d,** Simplified illustration of methionine cycle. Methionine, an essential sulfur-containing amino acid that is rich in animal products, enters the methionine cycle where it is converted to S-adenosylmethionine and subsequently to homocysteine, which can be remethylated back to methionine. Created using bioRender**. e,** Distribution of plasma homocysteine levels in PR individuals (n = 60) at both time points. **f,** Dosage effect of rs74891397 on plasma homocysteine levels in PR individuals (n = 60). Asterisks in panel **a, b, f** indicate significance as *: p.adjusted < 0.05; **: p.adjusted < 0.01; ***: p.adjusted < 0.001 (p.adjusted based on 1,000 permutations from steady-state cis-pQTL mapping); ns = non-significant. Asterisks in panels **c, e** indicate significance as *: p < 0.05; **: p < 0.01; ***: p < 0.001 (paired t-test); ns = non-significant. PR: periodically restricted group; NR: non-restricted group; T1: time point 1; T2: time point 2; p.adjusted = adjusted p-value; p = p-value.

When investigating plasma levels of methionine independently of genotype, we found a net marginal increase of its levels during animal product restriction (paired t-test; T2 vs T1; mean difference = 0.121; p = 9.05e-5) and a concurrent reduction in levels of homocysteine, a non-proteinogenic amino acid derived from, and recyclable back into, methionine^24^ (paired t-test; T2 vs T1; mean difference = −0.106; p = 0.001) (**Fig. 4c-e)**. Outside of the methionine cycle, homocysteine can also be converted to cysteine through the transsulfuration pathway, which plays a central role in cellular redox homeostasis^25^. However, diet-associated differences in cysteine levels were not found (paired t-test; T2 vs T1; mean difference = 0.012; p = 0.875) (**Supplementary** Fig. 2) suggesting conversion of homocysteine back to methionine to counterbalance dietary deficit of methionine during restriction. Further studies measuring biochemical intermediates in methionine metabolism however are necessary to elucidate effects of animal product restriction. Although rs74891397 showed diet-responsive regulation of MSRA and plasma methionine, its effects did not extend to homocysteine (p = 0.762 at T2) and cysteine (p = 0.327 at T2) indicating that its influence may be restricted to specific nodes of the methionine cycle (**Fig. 4e, Supplementary** Fig. 2).

### Seasonally regulated genetic effects on SERPINB1 associated with regulation of neutrophils

Loss of genetic regulatory effects on SERPINB1 abundance was detected for both dietary groups at T2 (rs2293772; lfsr = 0.008 in PR at T1; lfsr = 0.001 in NR at T1) (**Fig. 2**). SERPINB1 has a key role in neutrophil regulation^26^ acting as a gatekeeper of inflammation by restraining serine proteases and caspases^27^. We and others have shown that neutrophil numbers display strong seasonal fluctuation, with numbers decreasing during spring (T2)^10,28^. Genetic effects of rs2293772 extended to neutrophil counts in PR individuals at T2 (linear regression model; beta = −0.211; se = 0.103; p = 0.042 in PR at T2) (**Supplementary** Fig. 3), in whom reductions were more prominent. There is growing evidence that immune function exhibits seasonal variation^29^, which is also reflected in the seasonal patterns of various diseases, including cardiovascular disease, allergies, autoimmune disorders, and psychiatric conditions^30^. Our findings suggest genotype-dependent seasonal variation in immune system-related protein levels, which may contribute to individual differences in seasonal susceptibility to these diseases.

### Season-dependent genetic effects on apoptosis-related proteins may be masked by animal product restriction

In the control group, we detected gain of genetic regulatory effects for a single protein at T2 (MAVS) and three instances of loss of genetic regulatory effects (CASP3, PDLIM7, IL12RB1) (**Fig. 2**). MAVS is an outer mitochondrial membrane protein with a role in viral infection control through induction of caspase-dependent apoptosis^31^. CASP3, also located in mitochondria, is a key apoptosis effector protein, while PDLIM7 functions as a ubiquitin E3 ligase, inhibiting NF-κB-mediated inflammatory responses^32^. E3s are important regulators of mitochondrial and receptor-mediated apoptosis through their role in ubiquitination^33^. Finally, IL12RB1 is a receptor for IL-12^34^, an immunoregulatory cytokine involved in cell-mediated immunity which induces cell apoptosis. All four proteins are part of a protein-protein interaction network that is enriched for components of the death-inducing signaling complex (functional enrichment analysis; FDR = 2.96e-5), with MAVS, CASP3, and PDLIM7 being directly connected to TRADD, a mediator of programmed cell death and NF-κB activation^35^ (**Supplementary** Fig. 4). Based on these results, it is plausible that shared biological processes, stemming from seasonal changes in immune system function, may drive the gain or loss of proteogenomic regulatory effects. Given the impact of dietary restriction of animal products on the immune system^3,9,10,36^, these potential season-dependent genetic effects may be suppressed by animal product restriction and are thus not detected in the PR group.

Colocalization of differential cis-pQTLs with GWAS traits revealed that IL12RB1 differential cis-pQTL rs11668601 (**Fig. 5a**) colocalized with GGT (PPH4 = 0.978 in NR at T1) and with 11 additional traits, mostly reflecting liver function, only in NR at T1 (**Fig. 5b, Supplementary Table 5)**. rs11668601 is located 17 kb downstream of the IL12RB1 TSS and maps on a predicted binding site for transcription factors NR1D1/NR1D2. These transcription factors play an important role in diurnal immune function by regulating core circadian clock genes such BMAL1^37^. In turn, these signalling pathways accommodate seasonal variation in light duration, thus displaying patterns of seasonality^38^. Colocalization of CASP3 and PDLIM differential cis-pQTLs with intrinsic epigenetic age acceleration and arm fat percentage, respectively (**Extended Data Fig. 3 and 4**) suggests additional genotype-dependent seasonal effects on higher level traits. Further work addressing possible genotype-dependent seasonal effects, and whether these can be modified by diet, will enable us to understand the contribution of seasonality to disease risk.

**Figure 5.**
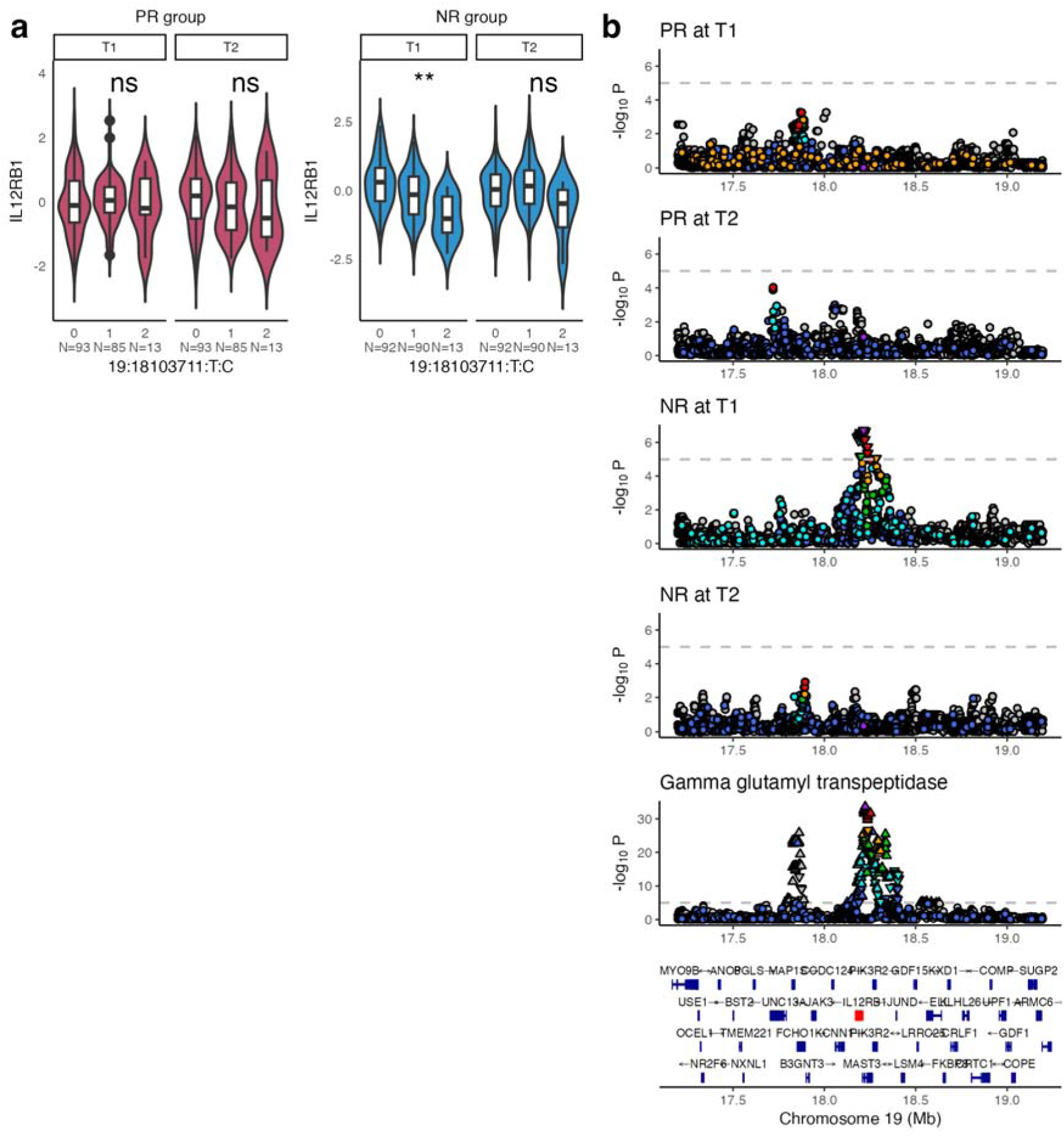
Differential cis-pQTL rs11668601 associated with IL12RB1 levels in the control (NR) group only at T1. **a,** Dosage effect of differential cis-pQTL 19:18103711:T:C (rs11668601) on IL12RB1 levels in PR and NR groups. **b,** Stacked regional plots for colocalization results of rs11668601 and GGT levels in each context. rs11668601 colocalizes with GGT levels, a biomarker of liver health in the NR group only at T1 (PPH4 = 0.98) and in no other context (PPH4 range: 0.04 to 0.08 in other contexts). Asterisks in panel **a** indicate significance as *: p.adjusted < 0.05; **: p.adjusted < 0.01; ***: p.adjusted < 0.001 (p.adjusted based on 1,000 permutations from steady-state cis-pQTL mapping); ns = non-significant. PR: periodically restricted group; NR: non-restricted group; T1: time point 1; T2: time point 2; p.adjusted = adjusted p-value.

### Regulatory variant for key metabolic regulator FGF21 colocalizes with eosinophil and platelet measurements

Our previous work has highlighted FGF21 as the protein most affected by animal product restriction (2.4-fold increased abundance)^9^. FGF21 is involved in regulation of energy homeostasis, lipid and glucose metabolism, and insulin sensitivity in humans and mice^39–41^. In mice, FGF21 promotes adipose tissue beiging and improved metabolic health^42^ and its prolonged overexpression has been shown to extend lifespan^43^. Although well-studied in model organisms, our knowledge on how this key metabolic hormone functions in humans lags behind^44^. While FGF21 cis-variants did not meet our stringent criteria for differential cis-pQTL assignment, we identified an FGF21-associated cis-pQTL only in the context of animal product restriction (rs4645881; p.adjusted = 0.022; q-value = 0.042 PR at T2) (**Fig. 6a**). rs4645881 showed evidence of colocalization with eosinophil counts (PPH = 0.92) and plateletcrit (PPH = 0.91) (**Fig. 6b, Extended Data Fig. 5**) and with basophil and lymphocyte counts (PPH4 = 0.75 and PPH4 = 0.66 respectively) (**Supplementary Table 7**). Studies in mice have shown that upon exposure to cold, adipose-derived FGF21 induces expression of CCL11 in mature adipocytes, which in turn promotes recruitment of eosinophils, M2 macrophage activation, and beiging of adipose tissue^45,46^. Although FGF21 has been shown to promote adipose tissue beiging in model organisms, its role in humans remains insufficiently characterised^47^. Our previous work has shown that plasma levels of CCL11 increase following animal product restriction and we have additionally predicted a role for this chemokine in human FGF21 signalling^9^. Although these findings should be interpreted with caution, it is plausible that diet-induced FGF21 levels may influence the process of adipose tissue beiging. We highlight the likely metabolic importance of eosinophils in humans and suggest that elucidating these signalling cascades may aid the identification of downstream effectors of FGF21 that could constitute potential therapeutics.

**Figure 6.**
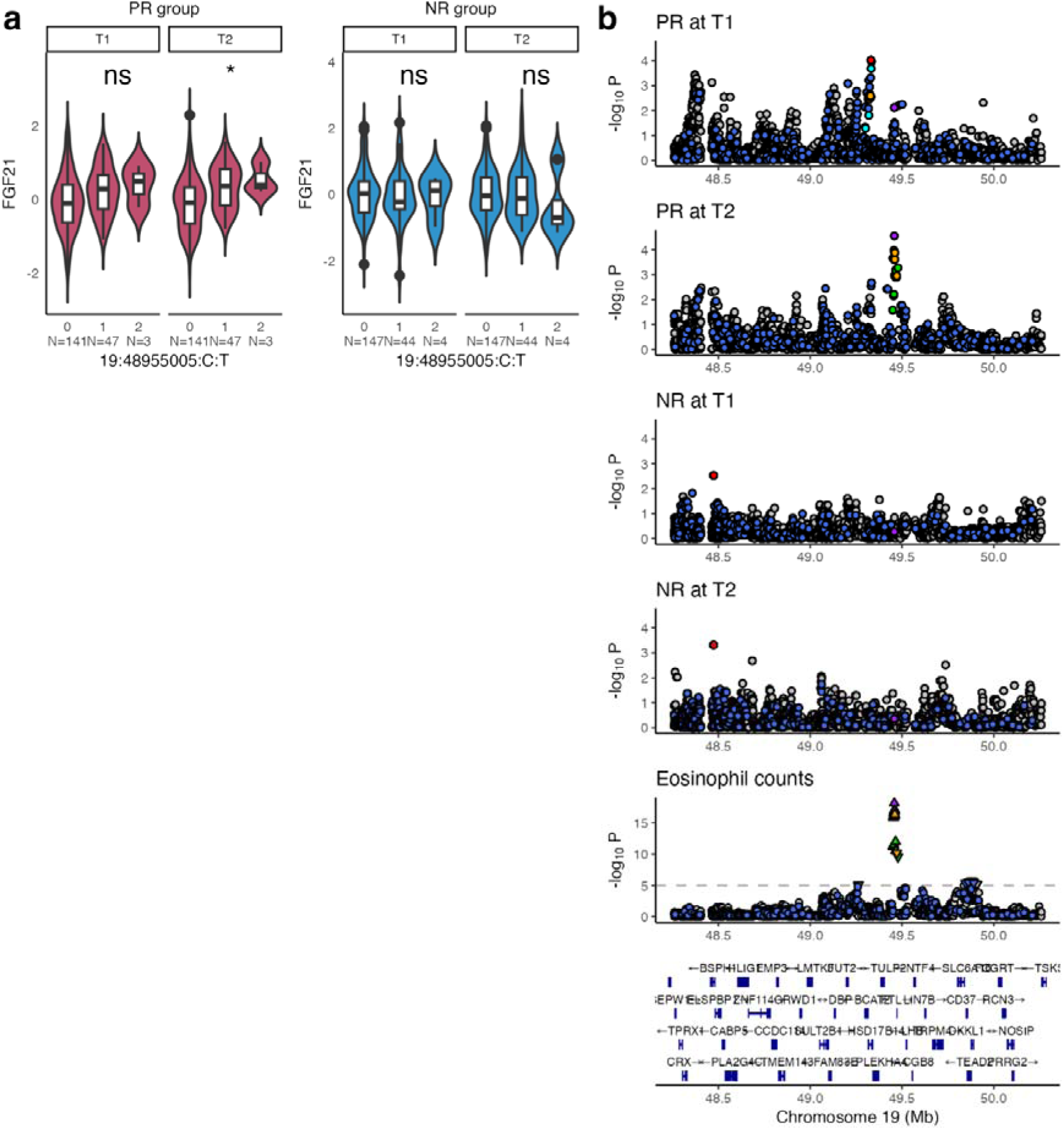
FGF21-associated cis-pQTL rs4645881 detected only during animal product restriction. **a,** Dosage effect of cis-pQTL 19:48955005:C:T (rs4645881) on FGF21 levels in PR and NR groups. **b,** Stacked regional plots for colocalization results of rs4645881 and eosinophil counts in each context. rs4645881 colocalizes with eosinophil counts only during dietary restriction (PPH4 = 0.92 in PR at T2) and in no other context (PPH4 range: 0.05 to 0.21 in other contexts). Asterisks in panel **a** indicate significance as *: p.adjusted < 0.05; **: p.adjusted < 0.01; ***: p.adjusted < 0.001 (p.adjusted based on 1,000 permutations from steady-state cis-pQTL mapping); ns = non-significant. PR: periodically restricted group; NR: non-restricted group; T1: time point 1; T2: time point 2; p.adjusted = adjusted p-value.

## Discussion

In the present study, we conducted proteogenomic analyses in apparently healthy individuals who alternate between omnivory and dietary restriction of animal products and demonstrated that dietary intake can drive differences in the cis-genetic regulation of plasma protein abundance in humans. While dietary interventions offer strong potential for disease management and prevention, the genetic factors influencing their effectiveness are still not well understood^7,48^. To our knowledge, this is the first study to demonstrate that short-term, real-world dietary patterns can modify genetically driven protein abundance in humans and to link these diet-responsive regulatory effects to broader health-related phenotypes.

We have revealed diet-responsive gain and loss of genetic regulatory effects on proteins that affect pathways of cellular homeostasis (LBR, MSRA, SNX9, ST6GAL1). LBR and MSRA in particular have roles in endogenous cholesterol synthesis and methionine metabolism respectively. We suggest that gain of genetic regulatory effects for these proteins occurs as a response to depleted dietary intake of cholesterol and methionine under animal product restriction. Diet-responsive genetic effects extended beyond LBR and MSRA regulation, with regulatory variants displaying associations with plasma levels of cholesterol-rich lipoproteins and methionine, respectively during animal product restriction.

We next showed that LBR diet-responsive cis-pQTL rs74148404 colocalizes suggestively with a variant linked to obesity. Although obesity is a complex trait shaped by multiple factors, including genetic and lifestyle parameters^17^, our findings suggest that there are genetic factors contributing to this trait, whose effects may be mediated under specific dietary conditions. This finding aligns with the observation that adherence to similar diets does not affect disease risk homogenously in a population^7^. Prospective studies in humans are needed to investigate how genetic variation interacts with diet to influence downstream clinical outcomes.

Similarly, we report a novel instance of diet-responsive gain of genetic regulatory effects on MSRA abundance. MSRA reduces oxidized methionine and rescues proteins damaged by oxidative stress. Dietary intake of methionine, an essential amino-acid, is substantially lower in vegan compared to omnivorous diets^22^. Moreover, reduced dietary intake of this amino acid has been associated with lower levels of oxidative stress, that are characteristic of vegan diets^22^, but the underlying mechanisms are not yet understood^25^. Restriction of methionine has also been associated with upregulation of liver-secreted metabolic regulator FGF21 in mice^39,49^. We found a cis-pQTL for FGF21, exerting its effects only under animal product restriction, which colocalized with eosinophil measurements. Studies in mice highlight a critical role for eosinophils in adipose tissue beiging^46^ while in humans adipose tissue eosinophil numbers have been shown to be lower in obese compared to non-obese individuals^45^. Our findings suggest that eosinophils may be a key modulator of adipose tissue beiging in humans, and that their action may be modulated by genetics, diet and their interaction.

Independently of dietary restriction, our work has revealed differences in proteogenomic regulation between time points for both dietary groups. These differential genetic regulatory effects may be driven by seasonality of immune system traits^29^ that is most likely a consequence of seasonal variation in pathogen exposure^30^. One of the most established seasonal effects is fluctuation of neutrophil abundance, with lower numbers present in spring compared to autumn^28^. Indeed, both dietary groups displayed loss of genetic regulatory effects of rs2293772 on levels of neutrophil regulatory protein SERPINB1. This suggests that seasonal effects may invoke genotype-dependent regulation of protein levels, and this interaction, in turn could contribute to effectiveness of immune response^29^.

Furthermore, we found differences in proteogenomic regulation between time points exclusively in the control group. These differential genetic regulatory effects were detected for immune system-related proteins within a network enriched for functions associated with programmed cell death (IL12RB1, MAVS, PDLIM7, CASP3). Apoptosis is a critical process in the function of the immune system^50^ and it may be involved in shaping seasonal variation of immune traits. While seasonality is experienced by both dietary groups, findings specific to the control group may reflect season by genotype effects that are masked by dietary restriction.

Our findings also revealed diet-responsive regulatory effects for proteins whose function is less well understood and merits further investigation. We uncovered diet-responsive loss of genetic regulatory effects on levels of SNX9, a member of the sorting nexin family, which when deleted in human *ex vivo* models, alleviates CD8+ T-cell exhaustion^51^. T-cell exhaustion can be induced by high cholesterol levels and targeting this protein has been proposed as a strategy to prevent exhaustion and enhance anti-tumor immunity^52^. Diet-dependent loss of SNX9 genetic regulatory effects may therefore be a response to reduced dietary intake of cholesterol. We also detected diet-dependent differences in the genetic regulation of protein-glycosylating enzyme ST6GAL1. In mice on a high-fat diet, ST6GAL1 expression is reduced in visceral adipose tissue, leading to enhanced adipogenesis and obesity^53^. Differences in genetic regulation during animal product restriction in this case may also be a response to reduced cholesterol intake. Given its potential involvement in pathways linked to cancer and atherosclerosis^54^, ST6GAL1 could be a promising candidate for further investigation.

Our work has several limitations. First, our study design enabled us to address dietary effects stemming from animal product restriction, but we have not addressed effects of dietary composition meaning that this study is limited to detecting changes driven by nutrients that are missing from the diet of participants. Second, protein abundance was measured for a limited number of proteins, capturing only a fraction of potential effects on protein abundance. Third, measurements were performed in plasma which reflects the biology of multiple tissues, not enabling us to address tissue-specific effects. Fourth, due to the limited number of IVs, we were unable to conduct MR analyses to address potential causal effects between proteins possessing differential cis-pQTLs and health outcomes. Finally, our work is limited to detecting effects of short-term dietary restriction, however longitudinal studies following these individuals over time will enable ascertainment of possible longer-term effects.

Investigation of proteogenomic regulation is key for explaining GWAS signals^13,55–57^. Investigating the proteome in longitudinal studies that capture dynamic effects across multiple time points will enhance our understanding of how regulatory variation responds to altered environmental conditions and how this affects disease risk. Overall, our findings suggest that dietary recommendations will not be equally effective across all individuals due to differential genetic effects. Considering how diet-responsive genetic regulatory effects can modulate disease risk will also be key for developing dietary interventions tailored to individual needs. Finally, the potential existence of genotype-dependent seasonal effects highlights a largely understudied field that may provide key insight in our understanding of mechanisms underlying health and disease.

## Methods

### Population sample and study design

A detailed description of the FastBio (religious <u>Fast</u>ing <u>Bio</u>logy) population sample has been outlined elsewhere^9,10^ (**Supplementary Table 1**). Briefly FastBio comprises two groups of individuals specified by their diet. Periodically restricted individuals (PR, N = 200) alternate between omnivory and abstinence from meat, fish, dairy products and eggs for 180-200 days annually, for religious reasons. This dietary pattern is initiated during childhood and is highly structured, involving four extended periods of animal product restriction throughout the year, as well as restriction on Wednesdays and Fridays. Typically, during animal product restriction, in addition to a pronounced reduction of animal fat intake, PR individuals undergo restriction of protein (as a proportion of total energy intake). This is driven by abstinence from almost all sources of animal protein, and is not accompanied by a reduction in total energy intake^11,12^. Non-restricted individuals (NR, N = 211) are a continuously omnivorous, control group. Both dietary groups were profiled at two time points: during a period of omnivory for both groups (T1) and following 3-4 weeks of animal product restriction for the PR group during Lent (T2). A 95% recall rate was achieved with final sample sizes of 192 and 195 individuals for the PR and NR groups respectively.

### Proteomic profiling

Proteomic profiling has been described in^9^. Briefly, 40 µL aliquot of plasma was taken forward to proteomic quantification on the Olink Explore 1536 platform (Olink Proteomics AB, Uppsala, Sweden). Protein levels for 1,472 analytes were reported in normalized protein expression values (NPX), which is a relative quantification unit in log2 scale. QC was performed for each sample and protein analyte, with QC-flagged values imputed using the protein’s mean NPX. To minimize the risk of bias, imputation was performed separately for each dietary group by time point combination (context). Proteins with more than 40% their values falling below the limit of detection (LOD) were excluded. The first two principal components were plotted with ±3SD boundaries, to identify outliers. Following QC, 1,218 proteins from 386 individuals were taken forward to analysis.

### Genotyping and imputation

Genomic DNA was extracted from blood using the iPrep PureLink gDNA Blood kit and iPrep Purification Instrument. Extracted DNA was genotyped on the Illumina Infinium GSA-23 v3.0. First degree relatives were excluded and variants with SNP call rate > 0.98 were retained, while duplicated variants were removed. Autosomal variants were imputed on Sanger Imputation Server using the Haplotype Reference Consortium (HRC.r1). The remaining SNPs were filtered for minor allele frequency (MAF) ≥ 0.05, info confidence score for the imputation quality INFO ≥ 0.8, and Hardy Weinberg Equilibrium (HWE) p-value > 1e - 6, resulting in a total of 5,120,744 genotyped and imputed autosomal SNPs for downstream analyses. Variants were initially in GRCh37 and were lifted over to GRCh38 build using https://genome.ucsc.edu/cgi-bin/hgLiftOver. To assess genetic ancestry of FastBio participants, we conducted principal component analysis (PCA) on 37,922 SNPs with MAF ≥ 0.05, alongside European reference populations from the 1000 Genomes project^58^ and from individuals of Turkish, Cypriot, Bulgarian and Ukrainian descent from^59^. The first two principal components (PC1, PC2) derived from the genotype data were plotted to visualize genetic ancestry of FastBio participants. SNP processing and filtering steps were performed using plink v1.90b6.16.

### Steady-state cis-pQTL mapping in each context

We performed cis-pQTL mapping for all four dietary group by time point combinations (contexts): PR at T1, PR at T2, NR at T1, NR at T2. Protein NPX values were first transformed using the inverse normal transformation formula (INT). We used TensorQTL^60^ to associate genotypes from 4,349,373 cis SNPs (within a 2 Mb window centred on the TSS of each gene), with levels of 1,181 proteins (excluding proteins on chromosome X) for each context, adjusting for: a) age, sex, BMI and medication use, b) context-defined protein principal components (PCs) from protein values, defined by the elbow method (9, 11, 10, 11 PCs for PR at T1, PR at T2, NR at T1, NR at T2 respectively), and c) the first three genotype PCs based on an LD-independent subset of SNPs using plink v2^61^ to correct for population structure. For each protein we performed 1,000 permutations to correct for multiple testing and we further adjusted results at FDR<0.05, using the qvalue method. Conditionally independent cis-pQTLs were mapped using the in-built function “-- mode cis_independent” of TensorQTL using a stepwise regression selection method as described in^62^. For each protein we also estimated the percentage of variance explained by the top cis-pQTL in each context as *Variance_explained_* = 2 · *MAF* · (1- *MAF*) · *b̂*^2^ ^63^.

### Replication of SNP-protein associations in UKB-PPP

We explored replication of the non-redundant union of all top cis-pQTLs across contexts (k = 630) by querying the same SNP-protein pairs in the UKB-PPP^13^. Publicly available summary statistics of UKB-PPP were downloaded via the synapse database (https://www.synapse.org/Synapse:syn51365303). Cis-pQTLs identified in FastBio that had a p-value lower than the Bonferroni corrected threshold of 7.94e-5 (0.05/630) in the UKB-PPP were considered to be replicated.

### Characteristics and annotation of cis-pQTLs

We annotated all independent cis-pQTLs across all contexts using VEP (https://www.ensembl.org/Tools/VEP) and explored cis-pQTL regulatory potential using RegulomeDB^64^.

### Differential cis-pQTL mapping

To identify shared and differential cis-pQTLs between contexts, we applied a multivariate adaptive shrinkage method using the mashr R package^65^ to jointly analyse SNP-protein associations derived from steady-state mapping across all contexts. Using the default workflow (https://stephenslab.github.io/mashr/articles/eQTL_outline.html) we distinguished cis-SNP-protein summary statistics into: a) a training set, consisting of the total number of tests (4,349,373) performed in the steady-state cis-pQTL mapping, and b) the most significant (lowest p-value) SNP-protein association from the four contexts, for each of the 1,181 proteins studied. We fitted the model to the training dataset to learn the mixture weights and scale the coefficients and subsequently extracted the posterior estimates for the strong signals from b). Significant cis-pQTLs were defined at a local false sign rate (lfsr) < 0.05. Using the posterior estimates we manually tested for SNP-protein pairs that were significant in at least one of the two time points (T1; T2) and displayed a change in magnitude of effect > 1.5 between time points, similar to Lin et al^8^. This approach enabled us to uncover variants that exhibited differential effects between time points within the same dietary group. Variants that were both top TensorQTL-defined cis-pQTLs, had a lfsr < 0.05 and a change in magnitude of effect greater than 1.5 (i.e. a differential effect between contexts) were considered as differential cis-pQTLs.

### Validation of differential genetic effects of differential cis-pQLTs on blood traits

We sought to validate the effects of differential cis-pQTLs for cases where it was possible to quantify molecular entities likely affected by differential effects of pGene action (rs74148404-LBR, rs74891397-MSRA, rs2293772-SERPINB1). To interrogate effects of rs74148404 on LBR activity, we queried plasma metabolite data from total FastBio participants (Rouskas, K. et al^9^). Abundance of plasma cholesterol classes (k = 57) were associated with rs74148404 genotypes using the same approach as in pQTL mapping, controlling for FDR at 5%. Similarly, to validate effects of rs74891397 on MSRA activity, we quantified plasma levels of methionine, homocysteine, and cysteine in a subset of 60 individuals (27 carriers of the rs74891397 effect allele, and 33 non-carriers matched for age, sex, BMI and medication use) at both time points. Abundance of these amino acids was quantified through UPLC-MS/MS and was associated with rs74891397 genotypes as above. To validate effects of rs2293772 on SERPINB1 activity, we queried serum clinical chemistry data from total FastBio participants from Eleni Loizidou et al.^10^ and associated neutrophil numbers with genotypes at rs2293772.

### Colocalization of differential cis-pQTLs with GWAS traits

We conducted PheWAS colocalization to link differential cis-pQTLs to GWAS traits. Each differential cis-pQTL with each LD proxy (r^2^ > 0.8) was queried against OpenGWAS using the ‘ieugwasr’ R package and the dedicated PheWAS function to find associated GWAS traits at p-value < 1e-5. For each associated trait we retrieved associations mapping ±1Mb from the TSS of the pGene associated with the differential cis-pQTL. Next, we used the Coloc R package^66^ to perform colocalization between each trait and differential cis-pQTL using default parameters. We deemed suggestive evidence of colocalization at PPH4 > 0.5 and evidence of colocalization at PPH4 > 0.8. This approach was also applied on FGF21-associated pQTL rs4645881.

## Acknowledgements

The authors are grateful to the FastBio study participants and to the Interbalkan Hospital Staff. We would also like to thank Dr Dimitrios Rouskas, Dr Pavlos Rouskas and Dr Loukas Kipouros for their invaluable help with sample collection. We acknowledge the technical support of Core Facility Metabolomics and Proteomics, and Core Facility Genomics at Helmholtz Munich and thank Dr Stefanie Hauck, Dr Agnese Petrera and Dr Harald Grallert for their help. We would also like to thank Dr Nikolaos Panousis and Dr Anders Mälarstig for helpful discussions. This work was funded by an ERC grant to Dr Antigone Dimas (FastBio – 716998). Dr Ozvan Bocher has received funding from the European Union’s Horizon 2020 research and innovation programme under Grant Agreement No 101017802 (OPTOMICS).

## Extended Data

**Extended Data Figure 1.**
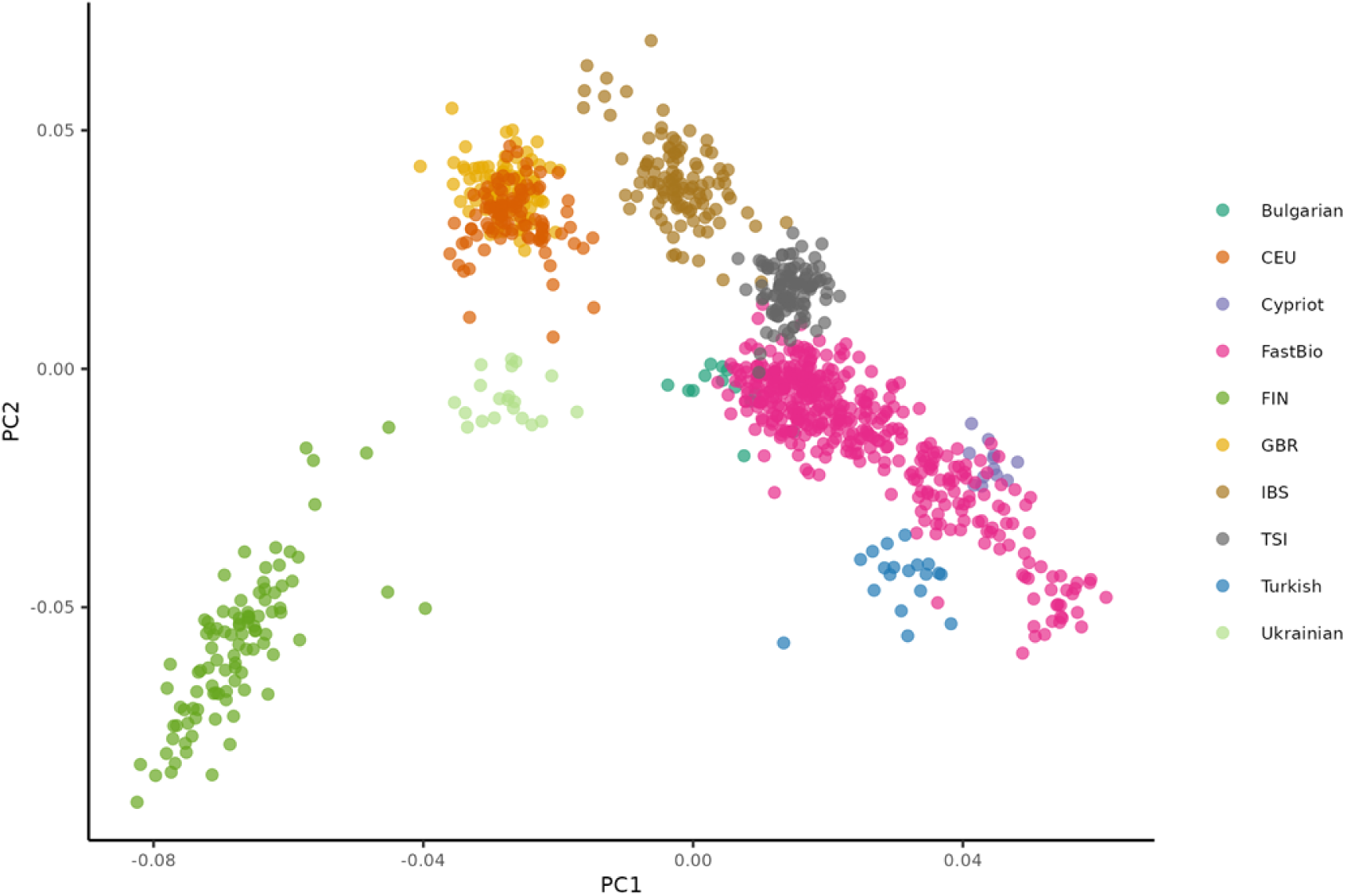
Principal component analysis (PCA) plot of genotypes from FastBio and reference populations. PCA plot depicting PC1, PC2 derived from genotype data from 37,922 SNPs. Colored data-points represent different populations. CEU, FIN, GBR, IBS and TSI populations are from the 1000 Genome project^58^, while Bulgarian, Cypriot, Turkish and Ukrainian individuals are from Bánfai, Z. et al^59^. Abbreviations: CEU, Utah residents (CEPH) with Northern and Western European ancestry; FIN, Finnish in Finland; GBR, British in England and Scotland; IBS, Iberian populations in Spain; TSI, Toscani in Italia.

**Extended Data Figure 2.**
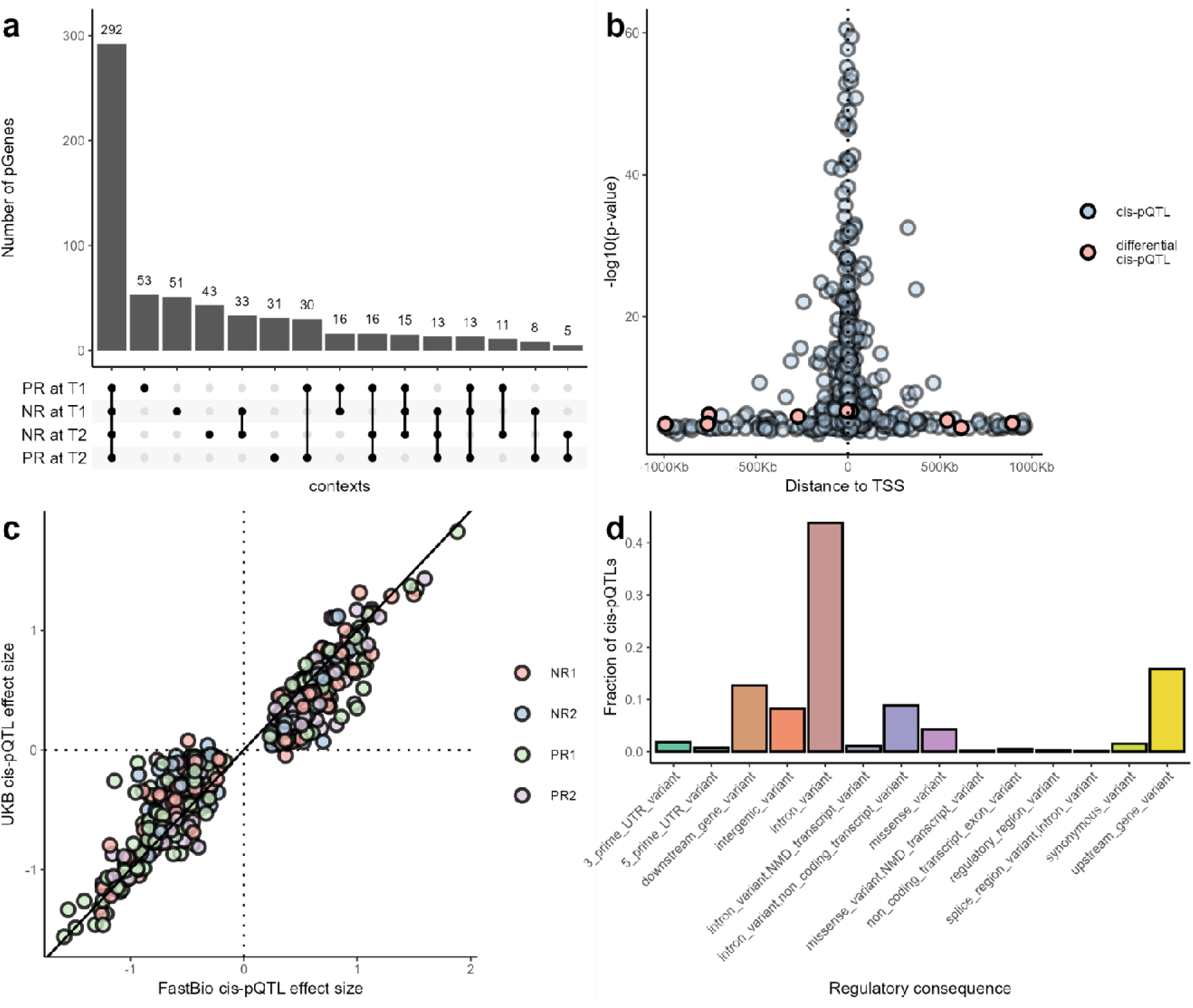
Steady-state cis-pQTL characteristics. **a,** Upset plot of steady-state cis-pQTLs detected in each time point by dietary group combination (context); each bar represents the number of proteins associated with at least one cis-pQTL (pGenes) (FDR < 0.05); connecting dots represent the intersections of pGenes between contexts. **b,** Distance to TSS of cis-pQTLs and significance of associations (-log10(p-value)); steady state cis-pQTLs shown in blue; differential cis-pQTLs shown in red. **c,** Direction and size of effect for top cis-pQTLs for each context in FastBio (x-axis), and in UKB-PPP (y-axis). **d,** Distribution of independent cis-pQTLs at each regulatory consequence (category) from Ensembl VEP.

**Extended Data Figure 3.**
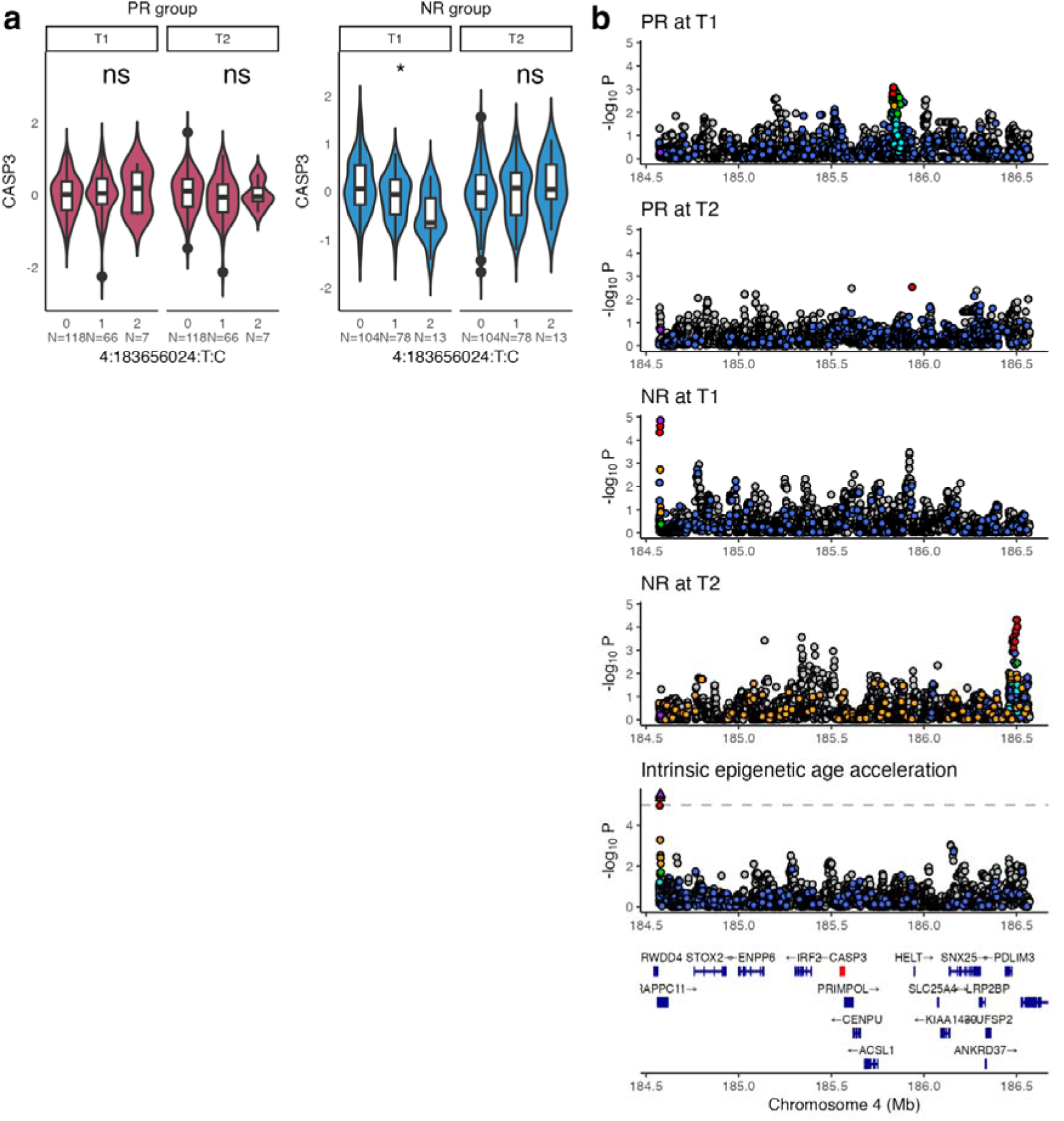
Differential cis-pQTL rs6838230 associated with CASP3 levels in the control (NR) group only at T1. **a,** Dosage effect of differential cis-pQTL 4:183656024:T:C (rs6838230) on CASP3 levels in PR and NR groups. **b,** Stacked regional plots for colocalization results of rs6838230 and intrinsic epigenetic age acceleration in each context. rs6838230 colocalizes with intrinsic epigenetic age acceleration in the NR group only at T1 (PPH4 = 0.97) and in no other context (PPH4 range: 0.02 to 0.04 in other contexts). Asterisks in panel **a** indicate significance as *: p.adjusted < 0.05; **: p.adjusted < 0.01; ***: p.adjusted < 0.001 (p.adjusted based on 1,000 permutations from steady-state cis-pQTL mapping); ns = non-significant. PR: periodically restricted group; NR: non-restricted group; T1: time point 1; T2: time point 2; p.adjusted = adjusted p-value.

**Extended Data Figure 4.**
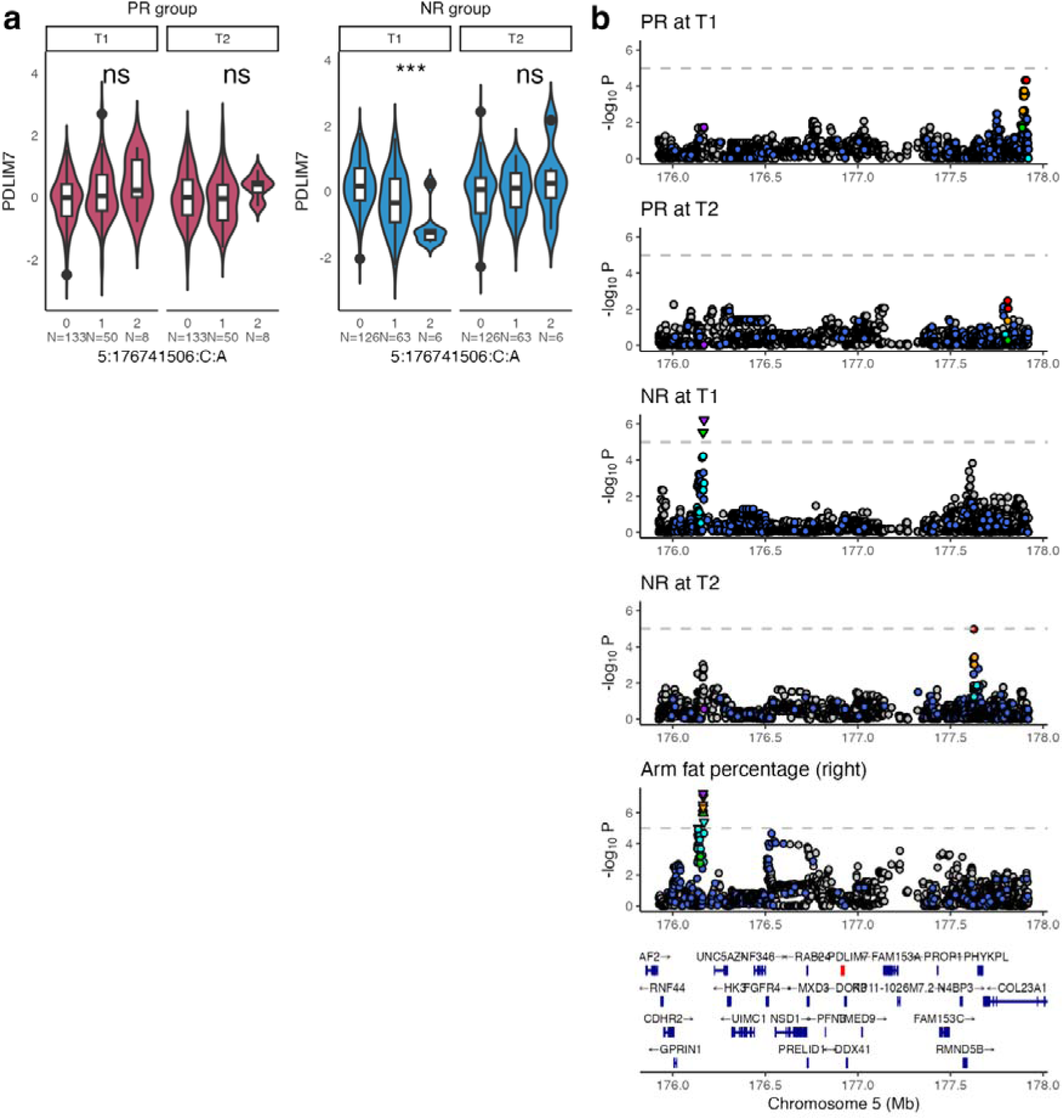
Differential cis-pQTL rs729823 associated with PDLIM7 levels in the control (NR) group only at T1. **a,** Dosage effect of differential cis-pQTL 5:176741506:C:A (rs729823) on PDLIM7 levels in PR and NR groups. **b,** Stacked regional plots for colocalization results of rs729823 and arm fat percentage (right) in each context. rs729823 colocalizes with arm fat percentage (right) in the NR group only at T1 (PPH4 = 0.94) and in no other context (PPH4 range: 0.04 to 0.35 in other contexts). Asterisks in panel **a** indicate significance as *: p.adjusted < 0.05; **: p.adjusted < 0.01; ***: p.adjusted < 0.001 (p.adjusted based on 1,000 permutations from steady-state cis-pQTL mapping); ns = non-significant. PR: periodically restricted group; NR: non-restricted group; T1: time point 1; T2: time point 2; p.adjusted = adjusted p-value.

**Extended Data Figure 5.**
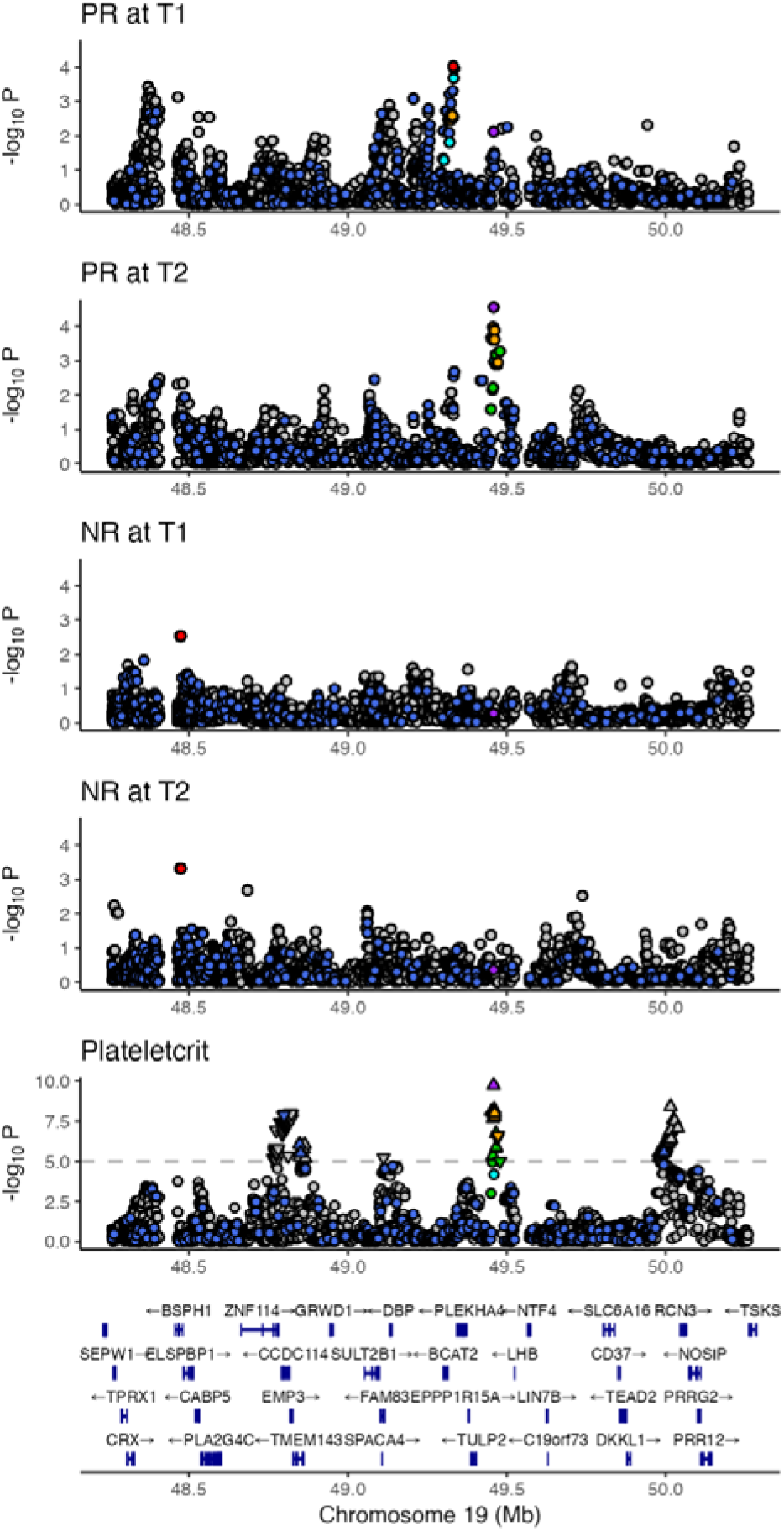
Regional colocalization plots for FGF21-associated cis-pQTL rs4645881 detected only during animal product restriction and plateletcrit. Stacked regional plots for colocalization results of 19:48955005:C:T (rs4645881) and plateletcrit in each context. rs4645881 colocalizes with plateletcrit only during dietary restriction in PR at T2 (PPH4 = 0.91) and in no other context (PPH4 range: 0.05 to 0.19 in other contexts).

**Extended Data Table 1.** Summary table of steady-state cis-pQTL mapping results. Context: diet by time point combination in which cis-pQTL mapping was performed. pGenes: proteins with at least one cis-pQTL. Cis-pQTLs: variants within 1Mb distance from the TSS of a gene that have been associated with the gene’s protein abundance (FDR < 0.05). Independent cis-pQTLs: cis-pQTLs that are conditionally independent in the tested cis-region.

| Context<br>(Sample size) | pGenes<br>(% of 1181 proteins) | Cis-pQTLs | Independent cis-pQTLs |
| --- | --- | --- | --- |
| PR at T1 (191) | 446 (38) | 41,806 | 585 |
| PR at T2 (191) | 408 (35) | 38,480 | 511 |
| NR at T1 (195) | 441 (37) | 39,674 | 559 |
| NR at T2 (195) | 428 (36) | 40,308 | 550 |

**Extended Data Table 2.** Summary table of colocalization results of differential cis-pQTLs with GWAS traits with at least suggestive evidence of colocalization (PPH4 > 0.5). Gene: protein coding genes associated with differential cis-pQTLs; no. colocalized traits: Number of traits with at least suggestive evidence of colocalization; top colocalizing trait: Trait with the highest probability of colocalization (PPH4); differential cis-pQTL: Differential cis-pQTL as candidate SNP for colocalization; top GWAS SNP: Most significant GWAS SNP in the specific region; LD (R^2^): R^2^ between differential cis-pQTL and top GWAS SNP in the tested region.

| Gene | no. colocalized traits | top colocalizing trait | differential cis-pQTL | top GWAS SNP | LD ($R^2$) |
| --- | --- | --- | --- | --- | --- |
| LBR | 1 | Diagnoses - secondary ICD10: E66.9 Obesity, unspecified | rs74148404 | rs34563765 | 0.35 |
| IL12RB1 | 42 | Gamma glutamyl transpeptidase | rs11668601 | rs273510 | 0.758 |
| FGF21 | 18 | eosinophil cell count | rs4645881 | rs4645881 | 1 |
| CASP3 | 1 | Intrinsic epigenetic age acceleration | rs6838230 | rs6818931 | 0.995 |
| PDLIM7 | 12 | Arm fat percentage (right) | rs729823 | rs729823 | 0.435 |

