## Supplementary Figures 1-4 & Supplementary Table 1. for "Diet-responsive proteogenomic effects following short-term restriction of animal products in humans"

**Supplementary Data**

|  | **Total (N=411)** | **PR (N=200)** | **NR (N=211)** | ***P*-value** |
| --- | --- | --- | --- | --- |
| Sex |  |  |  |  |
| *Female* | 224 (54.5) | 108 (54) | 116 (55) | 0.8425 |
| *Male* | 187 (45.5) | 92 (46) | 95 (45) |  |
| Age (yrs) | 48.1 ± 13.6 | 51.5 ± 13.5 | 45.0 ± 13.1 | <0.00001 |
| BMI (Kg/m^2^) | 27.3 ± 4.6 | 28.4 ± 4.6 | 26.2 ± 4.4 | <0.00001 |
| Blood Pressure (mmHg) |  |  |  |  |
| *Systolic BP (SBP)* | 124 ± 19.4 | 127 ± 19.0 | 121 ± 19.7 | 0.003 |
| *Diastolic BP (DBP)* | 79 ± 11.3 | 80 ± 10.8 | 78 ± 11.8 | 0.04 |
| Education |  |  |  |  |
| *Tertiary* | 293 (71.3) | 134 (67) | 159 (75.4) | 0.0612 |
| *Primary and Secondary* | 118 (28.7) | 66 (33) | 52 (24.6) |  |
| Marital status |  |  |  |  |
| *Married* | 286 (69.6) | 147 (73.5) | 139 (65.9) | 0.0931 |
| *Unmarried* | 125 (30.4) | 53 (26.5) | 72 (34.1) |  |
| Smoking (Y/N) |  |  |  |  |
| *Non-smokers* | 328 (79.8) | 187 (93.5) | 141 (66.8) | <0.00001 |
| *Smokers* | 83 (20.2) | 13 (6.5) | 70 (33.2) |  |
| Parental origin |  |  |  |  |
| *Northern Greece* | 275 (66.9) | 137 (68.5) | 138 (65.4) | 0.0979 |
| *Central or Southern Greece* | 32 (7.8) | 11 (5.5) | 21 (9.9) |  |

**Supplementary Table 1. Sociodemographic traits and geographic origin of study participants.** FastBio population sample has been described in detail in^1^. Continuous variables were expressed as mean ± standard deviations and categorical variables as N (%); SBP = Systolic Blood Pressure; DBP = Diastolic Blood Pressure; P-values are from chi-square test or Mann-Whitney test for categorical and numerical variables, respectively.

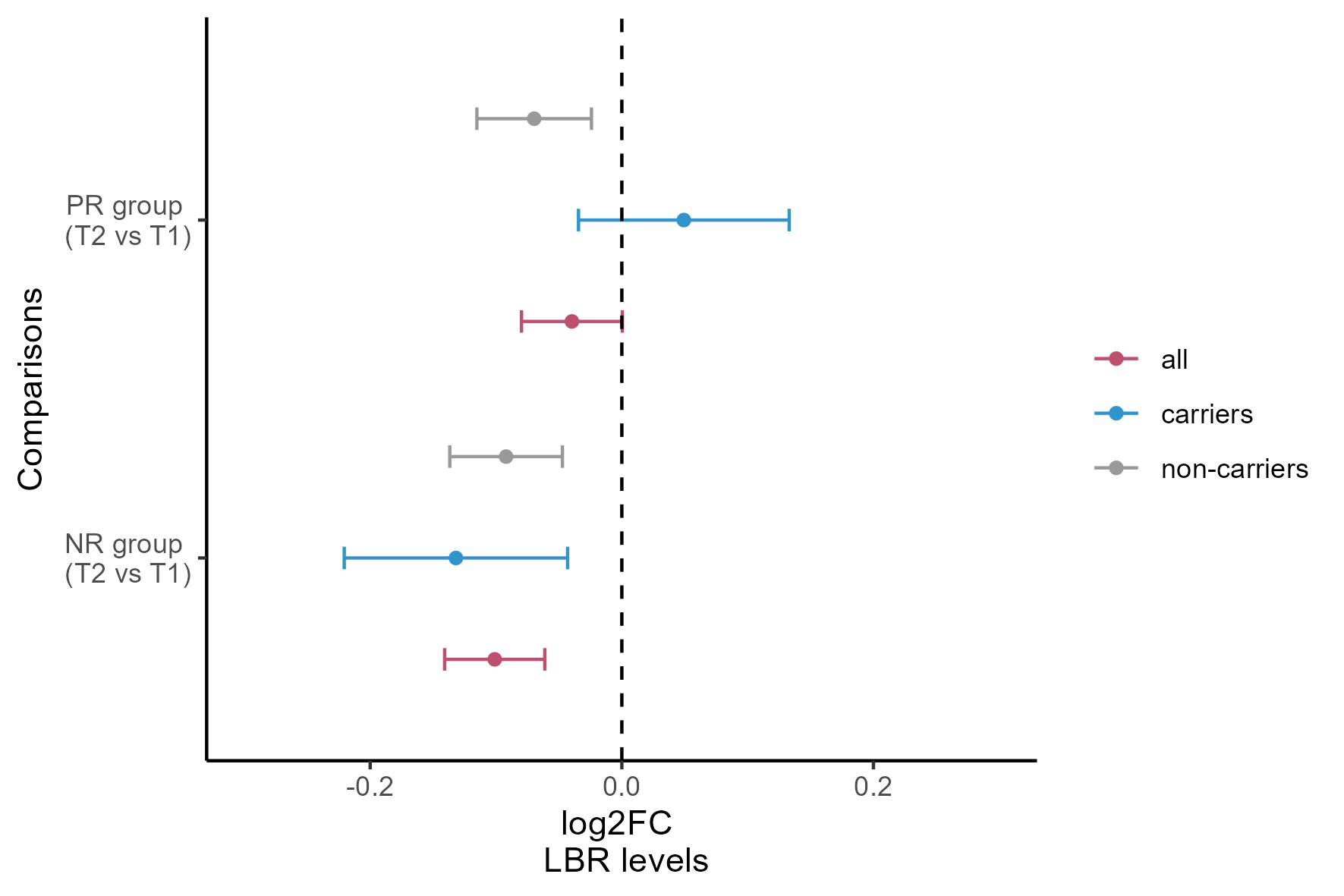

**Supplementary Figure 1. Differential abundance analysis of LBR stratified by diet-responsive C allele carriers at rs74148404**. Differential abundance analysis on LBR levels using limma model adjusting for mean protein levels, age^2^, sex, BMI and medication use. Point estimates represent the log2(FoldChange) (T2 vs T1) and the error bars the 95% confidence intervals. Color refers to the subset of individuals used in the analysis, red represents the reference analysis using all individuals (n = 385); grey represents a subset analysis excluding carriers of the differential cis-pQTL associated with LBR rs74148404 (n =293); blue represents the subset analysis including only carriers of the differential cis-pQTL associated with LBR rs74148404 (n = 92).

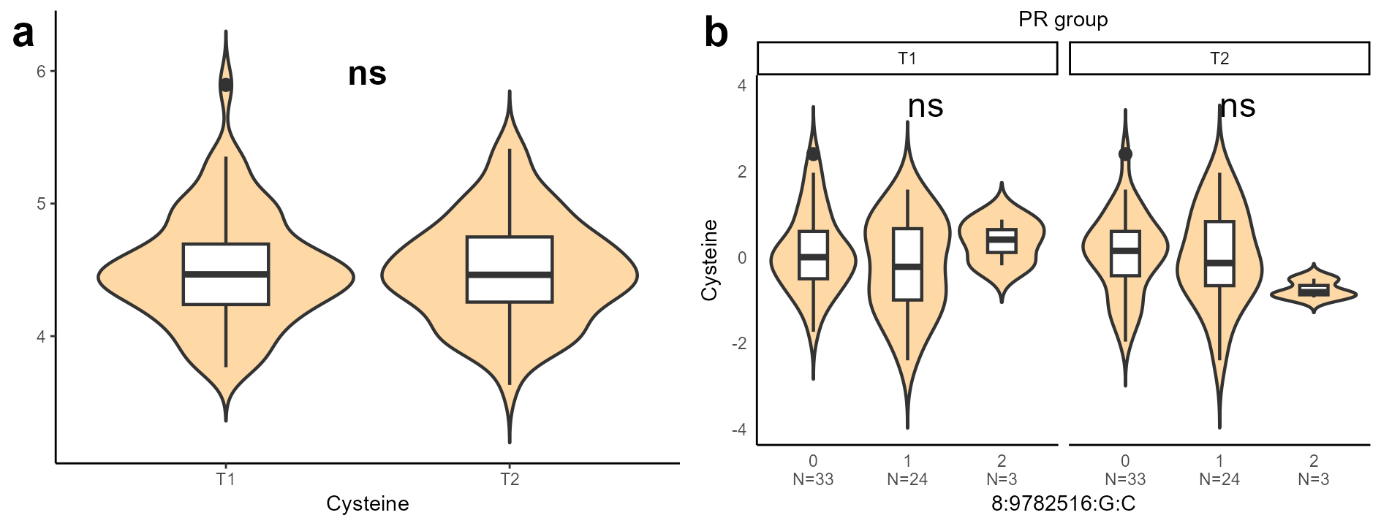

**Supplementary Figure 2. Distribution of cysteine levels in PR individuals and association testing with MSRA diet-responsive cis-pQTL.** Effects of dietary restriction did not extend to cysteine levels. **a,** Distribution of cysteine levels in PR individuals (n = 60) at both time points; **b,** Dosage effect of differential cis-pQTL 8:9782516:G:C (rs74891397) associated with MSRA on plasma cysteine residuals in PR individuals (n = 60). Asterisks in panel **a** indicate significance as *: p < 0.05; ** p < 0.01; *** p < 0.001 (paired t-test); ns = non-significant. Asterisks in panel **b** indicate significance as *: p < 0.05; **: p < 0.01; ***: p < 0.001 (linear model). PR: periodically restricted group; NR: non-restricted group; T1: time point 1; T2: time point 2; p = p-value.

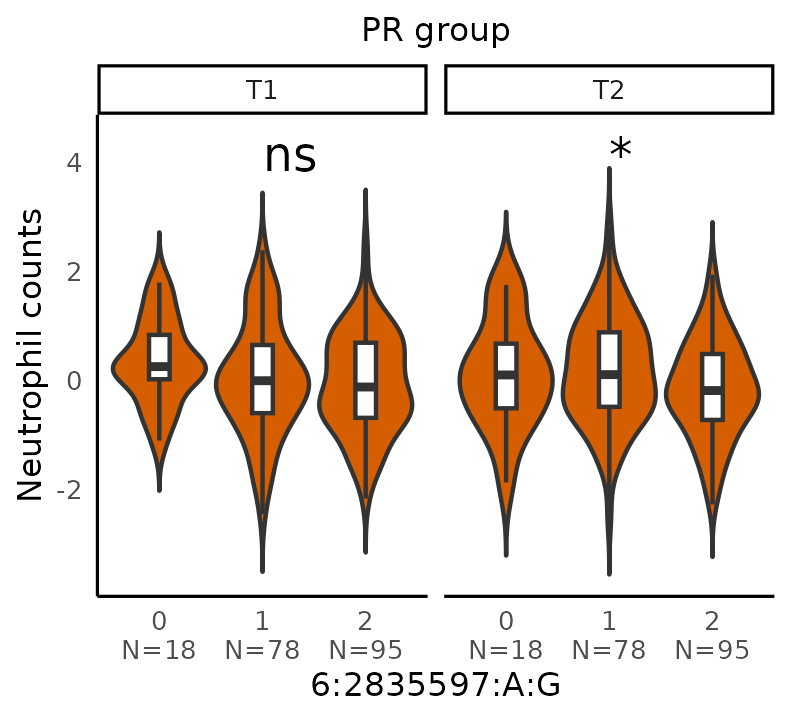

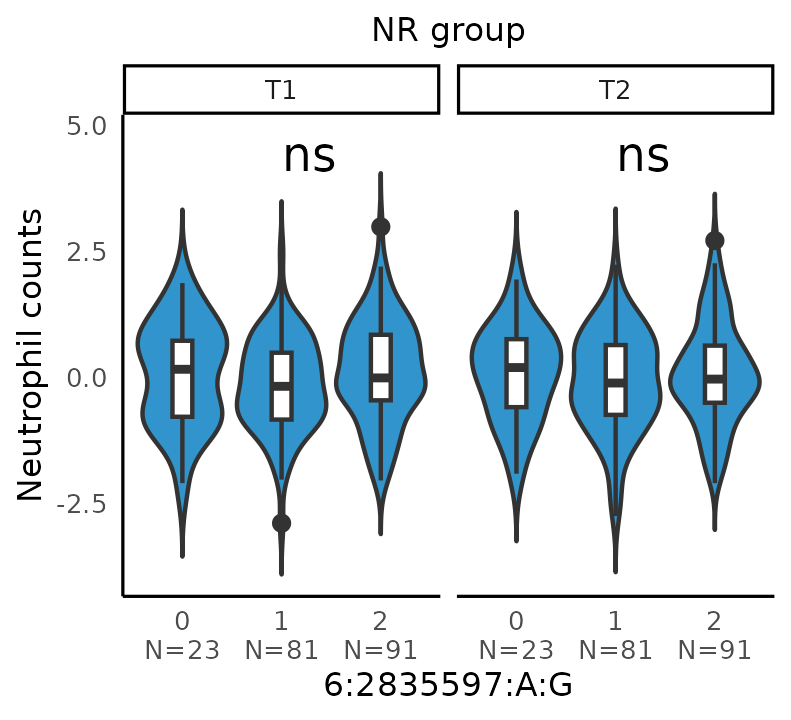

**Supplementary Figure 3. Association of SERPINB1 differential cis-pQTL rs2293772 with neutrophil counts**. Violin plot depicting the dosage effect of differential cis-pQTL 6:2835597:A:G (rs2293772) associated with SERPINB1 levels on the residuals of neutrophil counts in dietary group and time point. Asterisks indicate significance as *: p < 0.05; **: p < 0.01; ***: p < 0.001 (linear model); ns = non-significant. PR: periodically restricted group; NR: non-restricted group; T1: time point 1; T2: time point 2; p = p-value.

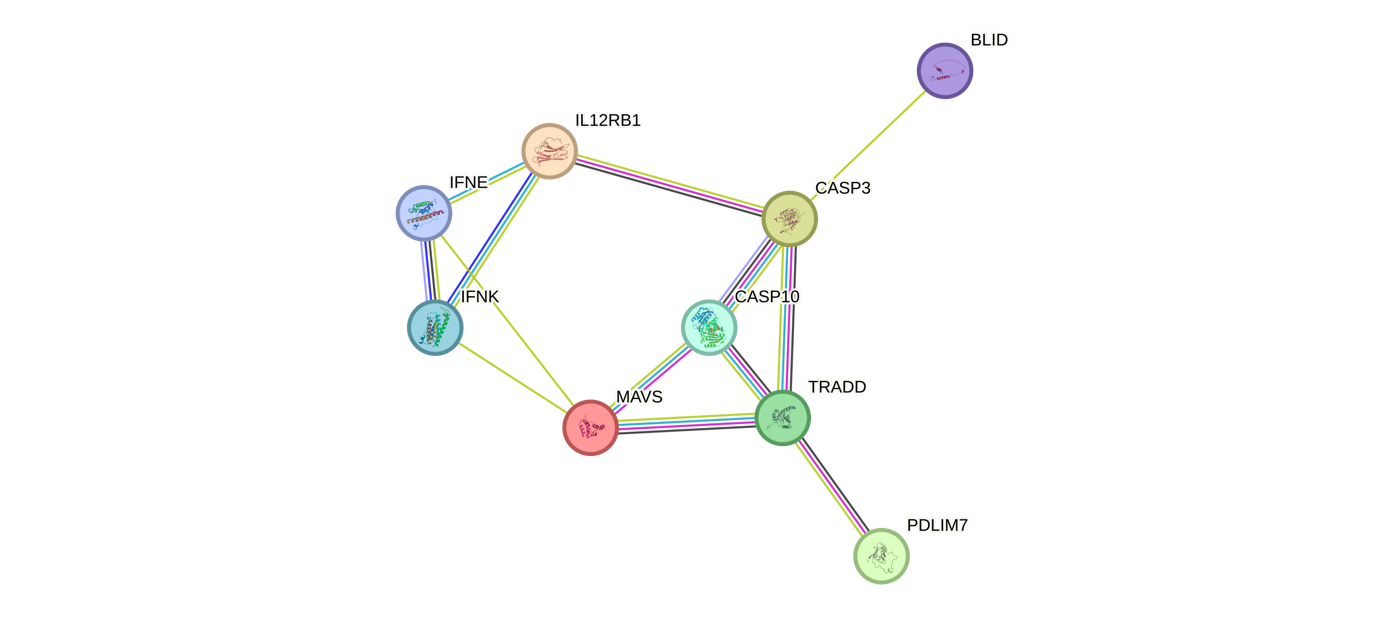

**Supplementary Figure 4.** **Protein-protein interaction network of proteins associated with differential cis-pQTLs unique to the control group**. The interaction network was produced by STRING (version 12.0) showing the connection of all proteins with differential cis-pQTLs unique to the control (NR) group (IL12RB1; MAVS; CASP3; PDLIM7).
